# Innate immune gene expression in *Acropora palmata* is consistent despite variance in yearly disease events

**DOI:** 10.1101/2020.01.20.912410

**Authors:** Benjamin Young, Xaymara M. Serrano, Stephanie Rosales, Margaret W. Miller, Dana Williams, Nikki Traylor-Knowles

## Abstract

Coral disease outbreaks are expected to increase in prevalence, frequency and severity due to climate change and other anthropogenic stressors. This is especially worrying for the Caribbean branching *Acropora palmata* which has already seen an 80% decrease in its coral cover, with this primarily due to disease. Despite the importance of this species, there has yet to be a characterization of its transcriptomic response to disease exposure. In this study we provide the first transcriptomic analysis of 12 *A. palmata* genotypes, and their symbiont Symbiodiniaceae, exposed to disease in 2016 and 2017. Year was the primary driver of sample variance for *A. palmata* and the Symbiodiniaceae. Lower expression of ribosomal genes in the coral, and higher expression of transmembrane ion transport genes in the Symbiodiniaceae indicate that the increased virulence in 2017 may have been due to a dysbiosis between the coral and Symbiodiniaceae. We also identified a conserved suite of innate immune genes responding to the disease challenge that was activated in both years. This included genes from the Toll-like receptor and lectin pathways, and antimicrobial peptides. Co-expression analysis identified a module positively correlated to disease exposure rich in innate immune genes, with D-amino acid oxidase, a gene implicated in phagocytosis and microbiome homeostasis, as the hub gene. The role of D-amino acid oxidase in coral immunity has not been characterized but holds potential as an important enzyme for responding to disease. Our results indicate that *A. palmata* mounts a similar immune response to disease exposure as other coral species previously studied, but with unique features that may be critical to the survival of this keystone Caribbean species.

## Introduction

Since the 1980’s, the Caribbean has seen dramatic losses of hard coral cover [1, 2]. This has been especially notable for *Acropora palmata* and *Acropora cervicornis*, which have seen an 80% reduction throughout their geographic range [2] resulting in them being classed threatened (US Endangered Species Act; ESA), and critically endangered (IUCN). The primary driver of this decline is disease [2–4] and this is particularly worrying for these species as climate change and anthropogenic stressors are now being implicated in increasing disease prevalence, frequency, and severity [5–10]. These two species are being heavily focused on for restoration activities in the Caribbean, but are historically susceptible to disease, thus it is imperative we understand the disease dynamics within the remnant populations.

Historically, coral disease research has focused on identifying the causative pathogens of coral disease with only a handful of studies fulfilling Koch’s postulates [11, 12]. This approach has proven difficult due to similar disease signs from coral species being attributed to different causative agents [13, 14], while shifting disease etiologies also causes disparity of causative agents over time [15, 16]. This is in part due to corals being symbiotic organisms that host a diverse set of microbial partners [17] and disentangling the roles of beneficial versus pathogenic is complex and will require interdisciplinary research efforts [11]. A new approach has been to use transcriptomics as a tool to understand the coral host’s genetic response to disease exposure and disease signs [18–25]. With the wide range of microbes that can potentially cause signs of disease, focusing on the host’s molecular ability to respond and resist infection has the potential to progress the coral disease field. Previous transcriptomic studies have led to the discovery that corals have a rich repertoire of putative innate immunity genes that are important in the response to disease exposure [26–29]. By focusing on understanding the host’s genes, it may be possible to characterize disease responses to a wide range of potential causative agents without definitively knowing exactly what they are. This will be particularly important in identifying signatures of disease resistance in coral species for restoration activities, while also providing potential diagnostic tools for coral health.

Despite both Caribbean Acroporid species being heavily incorporated into restoration practices, only the transcriptomic signature of *A. cervicornis* to disease exposure has been characterized [22, 23]. In this study we therefore provide the first transcriptomic analysis to disease exposure in *A. palmata*, as well as its symbiotic algal Symbiodiniaceae. The coral samples used in this study were previously tested to characterize genotypic patterns of resistance to disease grafting experiments run in 2016 and 2017 [30]. Little genotypic resistance was observed for *A. palmata* with all genotypes showing some replicates with transmission of disease signs over the course of the study [30]. There were differences in disease virulence, with 2017 (average 80% transmission) worse than 2016 (average 30% transmission). In this study we therefore focused on the transcriptomic response between healthy and disease outcome (disease vs no disease) rather than differences of resistance between genotypes. We hypothesized that there would be a clear transcriptomic disease response that was present in both 2016 and 2017 and that there would also be transcriptomic patterns which could explain differences in disease virulence between 2016 and 2017. We found that year showed the strongest correlation to overall gene expression for both *A. palmata* and the algal symbiont Symbiodiniaceae, with genes implicating a dysbiosis between host and symbiont behind the observed higher virulence in 2017. Response to disease exposure was only identified in *A. palmata*, with significantly differentially expressed genes involved in innate immune processes present. Coexpression analysis also identified two modules positively correlated to disease exposure, with this significantly enriched for lipid biosynthesis and innate immune processes [18–25].

## Methods

### Disease grafting experiment and genotype selection

For transcriptomic analysis, 12 *A. palmata* genotypes with previously published transmission information were analyzed [30]. In 2016 and 2017, disease grafting experiments were performed at the Coral Restoration Foundation (CRF; Key Largo Offshore Nursery) using 12 genotypes of *A. palmata* that are actively used for outplanting projects [30]. Using an isolated nursery structure, away from the main propagation nursery, fragments of *A. palmata* were grafted to diseased fragments of *A. cervicornis* over 7-days to identify disease transmission rates between the different genotypes. Reliable field disease diagnostics are lacking for most coral diseases including those affecting Caribbean Acroporids. Hence, disease inoculants were chosen according to gross visual signs and provide no guarantee that the disease etiology was the same between fragments and years [30]. At the base of the fragment ∼1cm^2^ piece of tissue was saved for nucleic acid extractions, these were taken before disease grafting (Baseline) and after 7-days exposure. After 7-days of exposure, fragment disease outcomes were scored as follows; Exposed: No Transmission (no visible disease signs, Fig 1B) or Exposed: Transmission (visible disease signs, Fig 1C). Samples were then either flash frozen in liquid nitrogen (2016), or placed in RNAlater (2017), and then stored at −80°C. In total for transcriptomic analysis, there were 32 samples in 2016 and 52 in 2017, with a breakdown of Baseline, Exposed: No Transmission and Exposed: Transmission shown in Table 1 and Fig 1A. Of the 12 total genotypes, three (HS1, ML6 and CN3) were assayed in both 2016 and 2017 (Table 1) to examine any impacts of each year on gene expression and ensure it was not due to genotypic variation.

**Fig 1.**
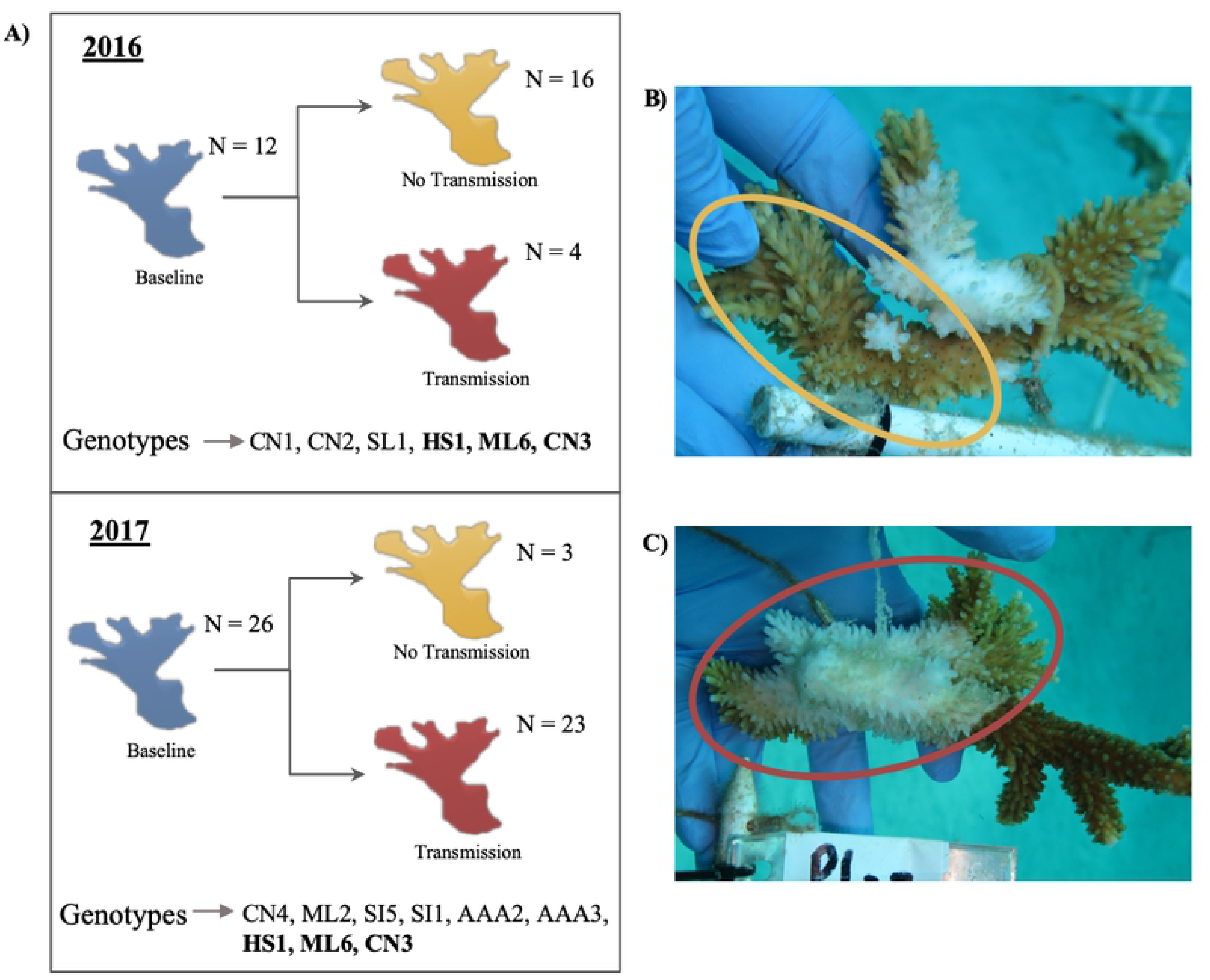
Experimental summary for transcriptomic analysis. A) A general overview of the field experiment conducted in 2016 and 2017 over 7-days each year. Samples were taken before grafting (Blue colony = Baseline) and after grafting, showing no signs of disease transmission (yellow colony = Exposed: No Transmission) or signs of disease transmission (red colony = Exposed: Transmission). Genotypes sequenced in each year are below colored coral fragments. B) The yellow circle indicates the apparently visually healthy *A. palmata* fragment grafted to the diseased *A. cervicornis* fragment after 7-days exposure. C) The red circle indicates *A. palmata* fragment showing disease signs grafted to the diseased *A. cervicornis* fragment after 7-days.

**Table 1:** Breakdown of genotypes and fragments sequenced for gene expression analysis.

| Genotype | Year | Baseline | Exposed: No Transmission | Exposed: Transmission | Total |
| --- | --- | --- | --- | --- | --- |
| CN1 | 2016 | 1 | 3 | 0 | 5 |
| CN2 | 2016 | 2 | 3 | 1 | 6 |
| SL | 2016 | 2 | 3 | 0 | 5 |
| HS1 | 2016 | 3 | 3 | 0 | 6 |
| ML6 | 2016 | 2 | 1 | 3 | 6 |
| CN3 | 2016 | 2 | 3 | 0 | 5 |
| HS1 | 2017 | 3 | 1 | 2 | 6 |
| ML6 | 2017 | 3 | 0 | 2 | 5 |
| CN3 | 2017 | 3 | 0 | 3 | 6 |
| CN4 | 2017 | 3 | 1 | 2 | 6 |
| ML2 | 2017 | 3 | 0 | 3 | 6 |
| SI5 | 2017 | 3 | 0 | 3 | 6 |
| SI1 | 2017 | 3 | 0 | 3 | 6 |
| AAA3 | 2017 | 3 | 0 | 3 | 6 |
| AAA2 | 2017 | 2 | 1 | 2 | 5 |

### cDNA library preparation and sequencing

A total of 88 samples were processed for total RNA extraction using the Qiagen RNeasy Minikit following the manufacturer’s protocol with the recommended 15-minute DNase digestion for all samples. Total RNA quality and quantity were assessed using a Nanodrop and Qubit fluorometer. Total RNA was then converted to complementary DNA (cDNA) libraries using Illumina TruSeq RNA Library poly A-tail selection prep kit following the manufacturer protocol. During cDNA library preparation, Illumina adaptors were randomly assigned to reduce bias between sequencing lanes. cDNA libraries were then quantified using a Qubit fluorometer and sent to the Utah Huntsman Cancer Institute High Throughput Genomics Shared Resource Center. cDNA quality control was performed using High Sensitivity D100 Screentape. A total of 84 samples passed quality control and were sequenced for 50 base pair single-end reads on 4-lanes using an Illumina HiSeq 2500.

### Bioinformatic analysis

Sequenced libraries were processed following standard practices for RNA-seq analysis [31]. All program parameters and scripts are available at (https://github.com/benyoung93/apal_disease_transcritpomics). Read quality was assessed using FastQC [32] and low-quality reads were trimmed using Trimmomatic [33]. Trimmed reads were then aligned to the *A. palmata* genome [34] using STAR [35] with the provided GFF file used for gene annotation and function. Because *A. palmata* shows stable symbioses with *Symbiodinium* (Clade A) over time and space [36], reads that did not align to the *A. palmata* genome were aligned to a *Symbiodinium* (Clade A) annotated transcriptome [37]. *A. palmata* and Symbiodiniaceae aligned reads where then quantified using Salmon [38] before being read into R (v3.6.1) and RStudio (v1.2.1335) using tximport [39]. An initial filtering for *A. palmata* (less than 1 count in greater than 15 samples), and for Symbiodiniaceae (less than 1 count in greater than 20 samples) was done using the counts per million (CPM) function in EdgeR [40]. Filtered counts were then used for differential gene expression analysis and co-expression analysis.

### Coral and Symbiodiniaceae principal components analysis

Sample counts were transformed using the variance stabilizing transformation (VST) function in DeSeq2 [41] and used as input for principal component analysis (PCA). A modified PlotPCA function was used to identify sample distribution for *A. palmata* and Symbiodiniaceae over multiple principal components (PCs) and plotted using ggplot2 [41]. To identify genes driving sample grouping in the PCA, loadings were extracted for PCs deemed interesting, and any genes with a +/- 2 standard deviation (SD) were retained for Gene Ontology (GO) analysis. We used a +/- 2 SD so to have a non-biased cut-off which was the same for each set of genes identified from *A. palmata* and Symbiodiniaceae.

### Coral host differential expression between Baseline and disease outcomes, and shared genes between contrasts

DeSeq2 [41] was used to analyze differential gene expression for the *A. palmata* quantified transcripts. The model ∼Year + Group was used to account for batch effects caused by different preservation methods used between the different years, while ‘Group’ encompassed Baseline and disease outcomes (Exposed: No transmission and Exposed: Transmission). This removed variance from the years and allowed significantly differentially expressed genes only due to disease outcome to be analyzed. Using this model, subsequent pairwise comparisons were performed using the contrast function in DeSeq2 between experimental outcomes; ‘Baseline VS Exposed: No Transmission’, and ‘Baseline Vs Exposed: Transmission’. Genes that were significantly differentially expressed (DEGs) had a false discovery rate (FDR) adjusted p value <0.01, and a Log 2-Fold Change (L2FC) >1 or <-1. These sets of DEG are used in GO analysis.

The two sets of significantly differentially expressed genes were then analyzed to identify any shared genes between the two contrasts (Baseline Vs Exposed: No Transmission, and Baseline Vs Exposed: Transmission). The L2FC for each contrast was compared to identify any differences in expression directionality due to disease outcome, and the full set of common genes are used in GO analysis.

### Weighted gene coexpression network analysis

To identify groups of coexpressed transcripts that correlated to Baseline and disease outcomes, a weighted gene coexpression network analysis (WGCNA; [44]) was used. Due to disease outcome being identified on PC axis 2 (Fig 2, B), the variance due to the year was removed using ‘removeBatchEffect’ in the program Limma [42]. Input data was therefore the CPM filtered batch removed counts with a VST for all 84 samples. Initial clustering using the Ward method in WGCNA [43] indicated there were no outlier samples and allowed retention of all 84 samples for coexpression analysis. A single signed network was built with manual network constructions (Key parameters: soft power = 12, minimum module size = 40, deep split = 2, merged cut height = 0.40, minimum verbose = 3, cutHeight = 0.997). The eigengene values of each module were correlated to disease outcome (Baseline, Exposed: No Transmission, Exposed: Transmission). To identify the highest connected gene within each module (hubgene), the WGCNA [43] command chooseTopHubInEachModule was used. All significant modules were then used in subsequent GO analysis.

**Fig 2.**
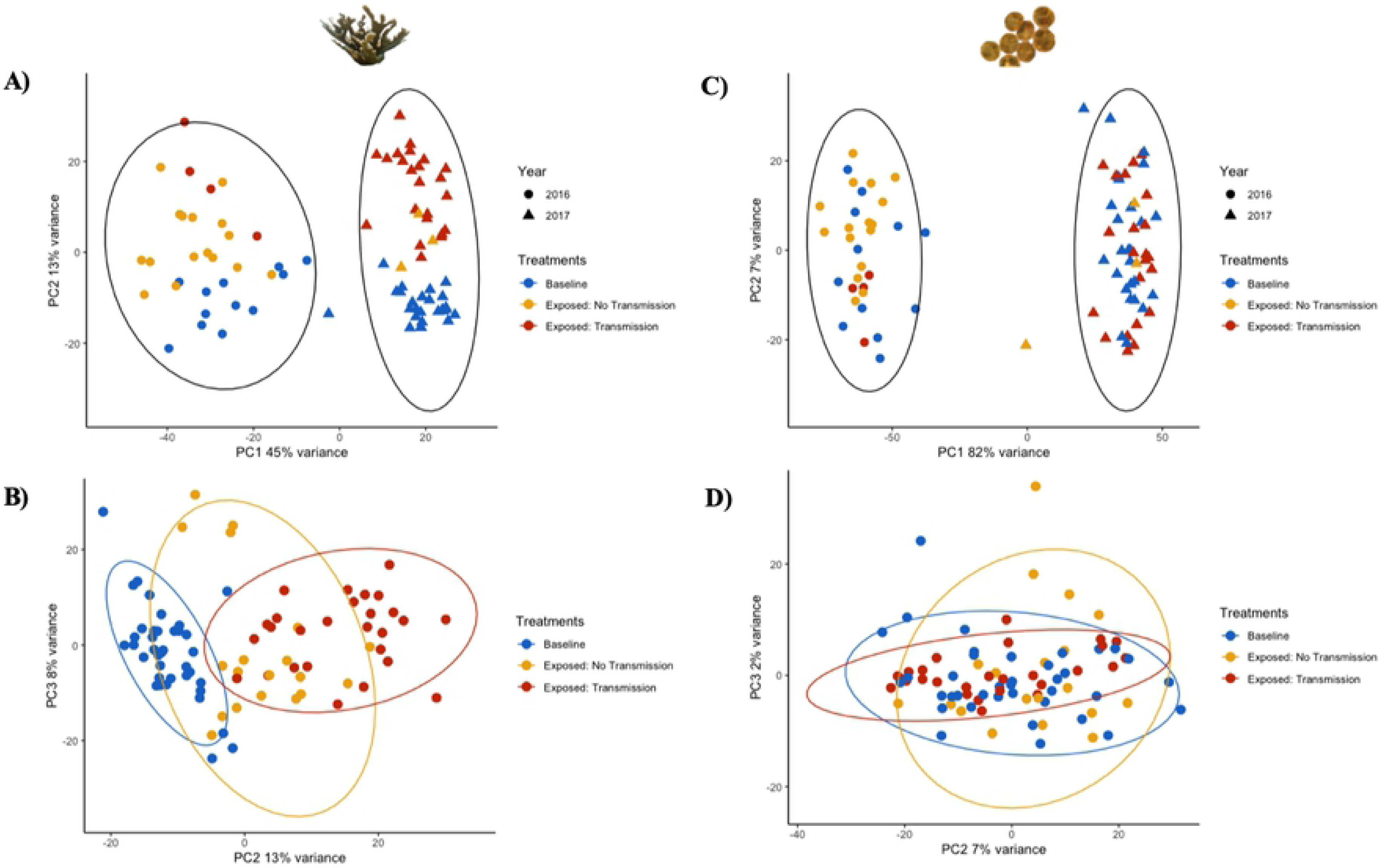
Coral and Symbiodiniaceae samples cluster firstly by year, while disease response is only identified in the coral. A) Principal Component (PC) 1 and PC2 of *A. palmata* counts, using a variance stabilizing transformation (VST), identifies the difference between years as the primary driver of sample variance. B) PC2 and PC3 of *A. palmata* counts, using a VST, is driven by disease outcome. C) PC1 and PC2 of Symbiodiniaceae counts, using a VST, identifies year as the primary driver of sample variance. D) PC2 and PC3 of Symbiodiniaceae counts, using a VST, shows no effect of disease outcome. For A) and C), black ellipses represent a 95% confidence interval in 2016 and 2017. For B) and D), the colored ellipses represent 95% confidence intervals for Baseline (blue), Exposed: No Transmission (yellow) and Exposed: Transmission (red).

### Gene ontology analysis

To identify significant enrichment of GO terms (biological process, cellular component, and molecular function) Cytoscape v3.7.2 [44], with the add-on application Bingo [45], was used. The hypergeometric test was utilized for GO enrichment and p-values were corrected with a Benjamini & Hochberg false discovery rate (FDR) correction (alpha set at < 0.01). The full mRNA transcriptome, available with the *A. palmata* genome [34], was used as the background set of genes for the enrichment tests. GO visualization was then done in Cytoscape v3.7.2 (45) allowing identification of significantly enriched relationships between parent and child terms. Genes in significantly enriched GO terms of interest were then visualized in RStudio using the VST counts and Complex Heatmap [46].

## Results

### Sequencing depth, read alignment, assignment metrics

A total of 84 samples were successfully sequenced on 4-lanes of an Illumina HiSeq 2500 with an average single-end read depth of 10,808,777. All raw reads are available on NCBI (SRA PRJNA529682). From quality filtered sequences, 74.64% of single end reads mapped to the *A. palmata* genome [34] using STAR [35]. Quantification, using Salmon [38], resulted in 35,079 genes having at least one count across all samples, with subsequent CPM filtering (less than 1 count in > 15 samples) reducing this to 18,913 genes for downstream analysis. Of reads not aligning to the *A. palmata* genome, an average of 21.54% aligned to the *Symbiodinium* (Clade A) reference transcriptome [37] using STAR [35]. Quantification using Salmon [38] yielded counts for 72,152 transcripts, with 28,035 of these retained for downstream analysis after CPM filtering (less than 1 count in greater than 20 samples).

### Year was the greatest driver of gene expression for *A. palmata* and Symbiodiniaceae gene expression with ribosomal and ion transport genes driving sample clustering

PCA showed *A. palmata* samples clustered by year on PC 1 (PC1 = 45%; Fig 2, A), followed by disease outcome on PC 2 (PC2 = 13%; Fig 2, B). Symbiodiniaceae samples also clustered by year on PC1 (PC1= 82%; Fig 2, C) while PC 2 showed no correlations to disease exposure or genotype (Fig 2, D).

Analysis of the genes driving PC1 variance for *A. palmata* identified 86 significantly enriched GO processes; 48 Biological Process, 6 Molecular Function, and 32 Cellular Components. Within Biological Process and Cellular Component, genes associated with ribosomal structure and function, as well as ribosomal RNA processing were significantly enriched. Three GO terms were also linked to immune processes; cell-cell adhesion, extracellular vesicular exosome, and apolipoprotein binding. Visualization of the VST counts for the genes within these GO terms identified 4 heatmap clusters (Fig 3, A). All genes linked to ribosomal processes showed lower normalized counts in 2017 than 2016, while GO terms with potential immune genes and functions showed higher normalized counts in 2017 than in 2016 (Fig 3, A). Principal component 1 loadings and full GO results for *A. palmata* are available in Supp 1.

**Fig 3:**
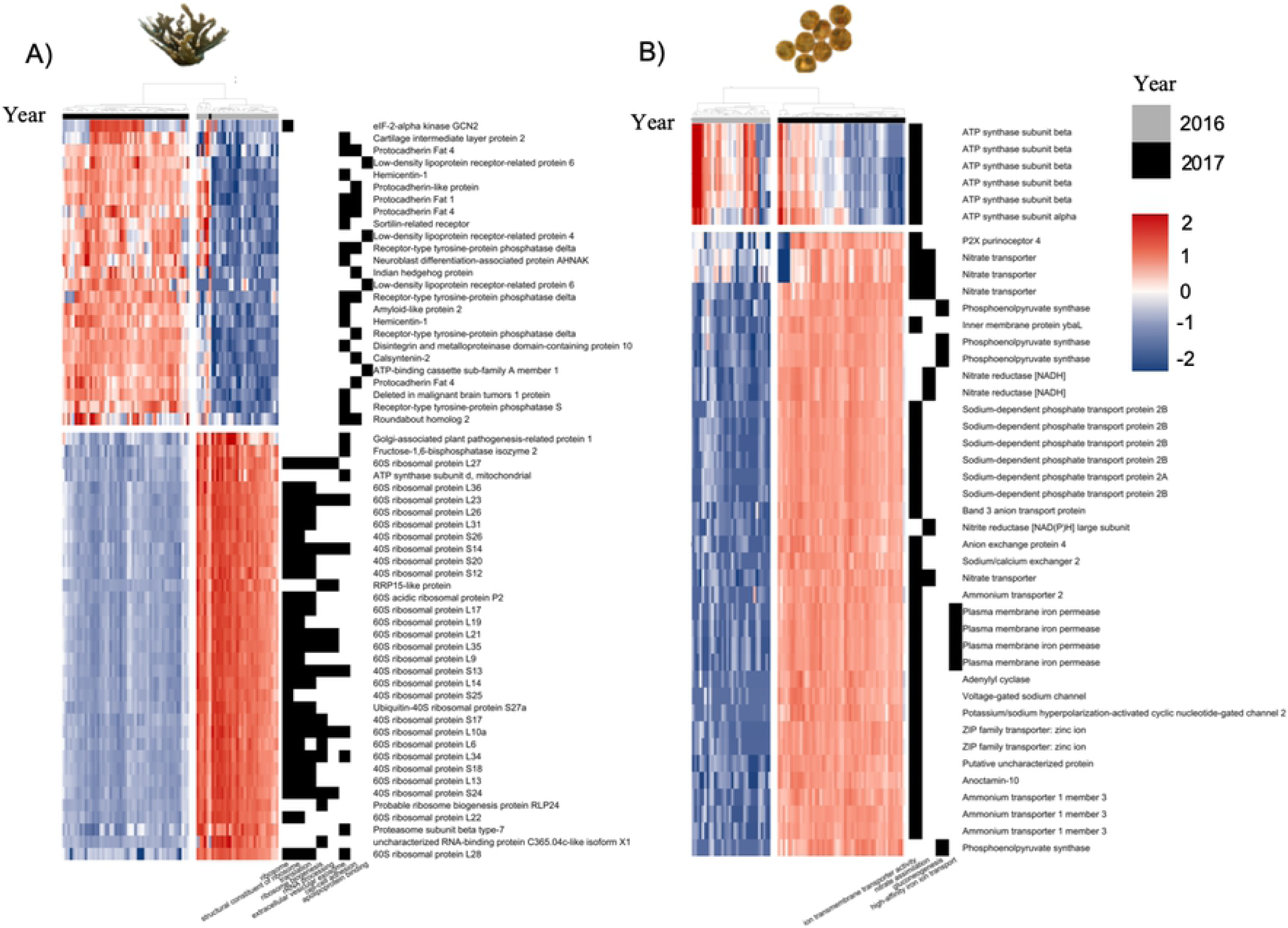
Genes driving the difference between 2016 and 2017 responses in the coral host and Symbiodiniaceae. A) Coral host genes linked to significantly enriched gene ontology (GO) terms, identified from principal component (PC) 1 loadings. Genes are linked to translation and ribosomal formation processes. Hierarchical clustering of the samples (heatmap columns) shows grouping between the samples from 2016 (grey) and 2017 (black), with 2016 genes having higher normalized expression and 2017 having lower normalized expression. B) Symbiodiniaceae genes linked to significantly enriched GO terms identified from PC1 loadings. Genes are linked to transmembrane ion transport processes. Hierarchical clustering of the samples (heatmap columns) shows grouping between the samples from 2016 (grey) and 2017 (black). For A) and B), grey = 2016 samples, black = 2017 samples. Left heatmap fill shows higher (red) to low (blue) gene counts using a variance stabilizing transformation. Right heatmap is presence (black) and absence (white) of genes to GO terms. Column dendrogram shows hierarchical clustering of samples. Rows (genes) also arranged using hierarchical clustering with dendrogram omitted.

For Symbiodiniaceae, there were 120 significantly enriched GO processes; 48 Biological Process, 6 Molecular Function, and 32 Cellular Components. In all three GO components, significantly enriched terms identified 2 main gene processes. Genes implicated in the transport of ions between cells and cellular components showed higher expression in 2017 than in 2016 (Fig 3, B). This included plasma membrane iron permease, nitrate and nitrite transporters, sodium transporters, zinc transporters, and ammonium transporters. Genes linked to photosynthesis, namely photosystems I and II in the light dependent reaction, also showed significant GO enrichment. The genes within these photosynthesis terms did not exhibit higher or lower expression compared between year, but instead showed a range of expression across the samples for each year (Supp 2). Principal component 1 loadings and full GO results for Symbiodiniaceae are available in Supp 3.

### Significant differential gene expression was identified between different disease outcomes in *A. palmata*

Differential gene expression analysis was only done for *A. palmata* due to there being no disease response identified in the Symbiodiniaceae. For Baseline Vs Exposed: No Transmission, there were 139 transcripts significantly downregulated, and 679 transcripts significantly upregulated, while Baseline Vs Exposed: Transmission had 678 transcripts significantly downregulated and 673 transcripts significantly upregulated (Fig 4A). Full lists of significant DEG for each contrast are available in Supp 4 and Supp 5 respectively. Between each contrast, there were 422 shared differentially expressed transcripts (Fig 4A). Of these, only 2 showed opposite LFC directionalities; a ‘PREDICTED cyclin-dependent kinase 11B-like partial’ (Baseline Vs Exposed : No Transmission L2FC = 2.57, Baseline Vs Exposed : Transmission L2FC = −1.98), and a Aspartate 1-decarboxylase (Baseline Vs Exposed: No Transmission L2FC = 1.53, Baseline Vs Exposed: Transmission L2FC = −2.18). A full list of shared genes with LFC is available in Supp 6.

**Fig 4:**
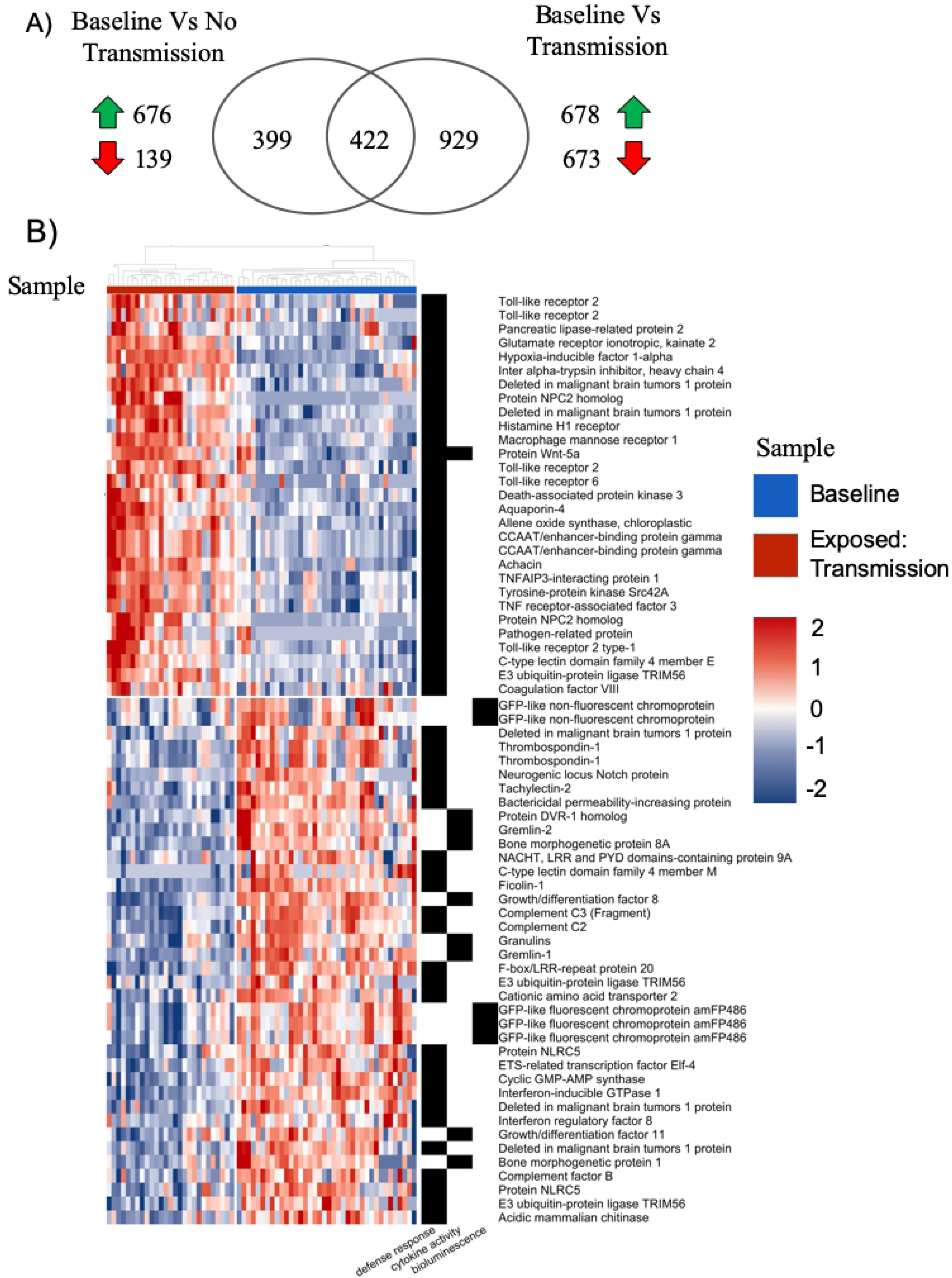
Unique and common genes between differential expression contrasts and significant innate immune genes in diseased corals. A) Venn diagram of the unique (left and right) and shared (intersect) differentially expressed genes from the two contrast arguments run in DeSeq2. The green arrow shows significantly upregulated, the red arrow shows significantly downregulated genes. B) Heatmaps showing genes linked to significantly enriched innate immune gene ontology (GO) terms identified from the Baseline versus Exposed: Transmission DeSeq2 contrast. Samples included are Baseline (blue) and Exposed: Transmission (red). Left heatmap fill shows higher (red) to low (blue) gene counts using a variance stabilizing transformation. Right heat map identifies genes present (black) or absent (white) from significantly enriched GO terms linked to innate immune response. Column dendrogram shows hierarchical clustering of samples. Rows (genes) also arranged using hierarchical clustering with dendrogram omitted.

#### Contrast between Baseline and Exposed: No Transmission

The significant DEGs for the contrast between Baseline and Exposed: No transmission showed significant enrichment of 18 GO terms (4 Biological Process, 7 Molecular Functions, and 7 Cellular Components). Biological Processes identified terms associated with cell adhesion and cell surface receptor linked signaling pathways including a number of putative immune function genes such as: tumor necrosis factors (TNFs), WNT proteins, protein kinase C epsilon type, and genes involved recognition such as Apolipophorin and C-type lectins. All significant GO terms and associated genes are available in Supp 4.

#### Contrast between Baseline and Exposed: Transmission

The significant DEGs for the contrast between Baseline and Exposed: Transmission showed significant enrichment of 46 Biological processes, 14 Cellular Component, and 35 Molecular Function. GO terms linked to Defense Response, Bioluminescence, and Cytokine Activity contained innate immune genes important in the main processes of innate immunity; recognition, signaling, and effector responses. Within these enriched GO terms there were a number of recognition innate immune genes, including four genes similar to Toll-like receptor (TLR) 2, and 2 genes similar to TLR 6 complexes (Fig 4, B). There were also lectin pathway recognition genes such as: C-type lectin domain family 4 member E and M, Ficolin-1. As well as other receptors which have been implicated in innate immunity; F-box/LRR-repeat protein 20, Histamine H1 receptor, Macrophage mannose receptor 1, two NOD-like receptor proteins, and a neurogenic locus notch protein (Fig 4. B). Innate immune genes involved in signaling pathways were also present, including TLR signaling pathway components such as; Deleted in malignant brain tumour 1, CCAAT/enhancer-binding protein gamma, Gremlin 1 and 2, NACHT LRR and PYD domain contain proteins 12 and 9A, TNF receptor-associated factor 3, TNFAIP3-interacting protein 1, and E3 ubiquitin-protein ligase TRIM56 (Fig 4, B). There were also genes important in lectin signaling; complement C2 and C3 fragments. Finally, there were genes involved in effector responses of innate immunity including antimicrobial peptides (AMPS) such as Achacin and Bactericidal permeability-increasing protein, and a pathogen related protein. Two genes identified as transcription factors; CCAAT/enhancer-binding protein gamma, and Interferon-inducible GTPase 1 and interferon regulatory factor 8 (Figure 4B). All significant GO terms and associated genes are available in Supp 5.

### Co-expression analysis identifies positively correlated modules of immune genes and lipid biosynthetic processes to disease exposure

After merging of similar modules, we identified 19 coexpressed modules that contained 76 to 2027 genes (Fig 5A, Supp 7). Of these 19 modules, 8 showed significant correlations to Baseline and disease outcomes (Exposed: No Transmission, and Exposed: Transmission) (Fig 5 B). Gene lists for all modules is provided in Supp 8.

**Fig 5:**
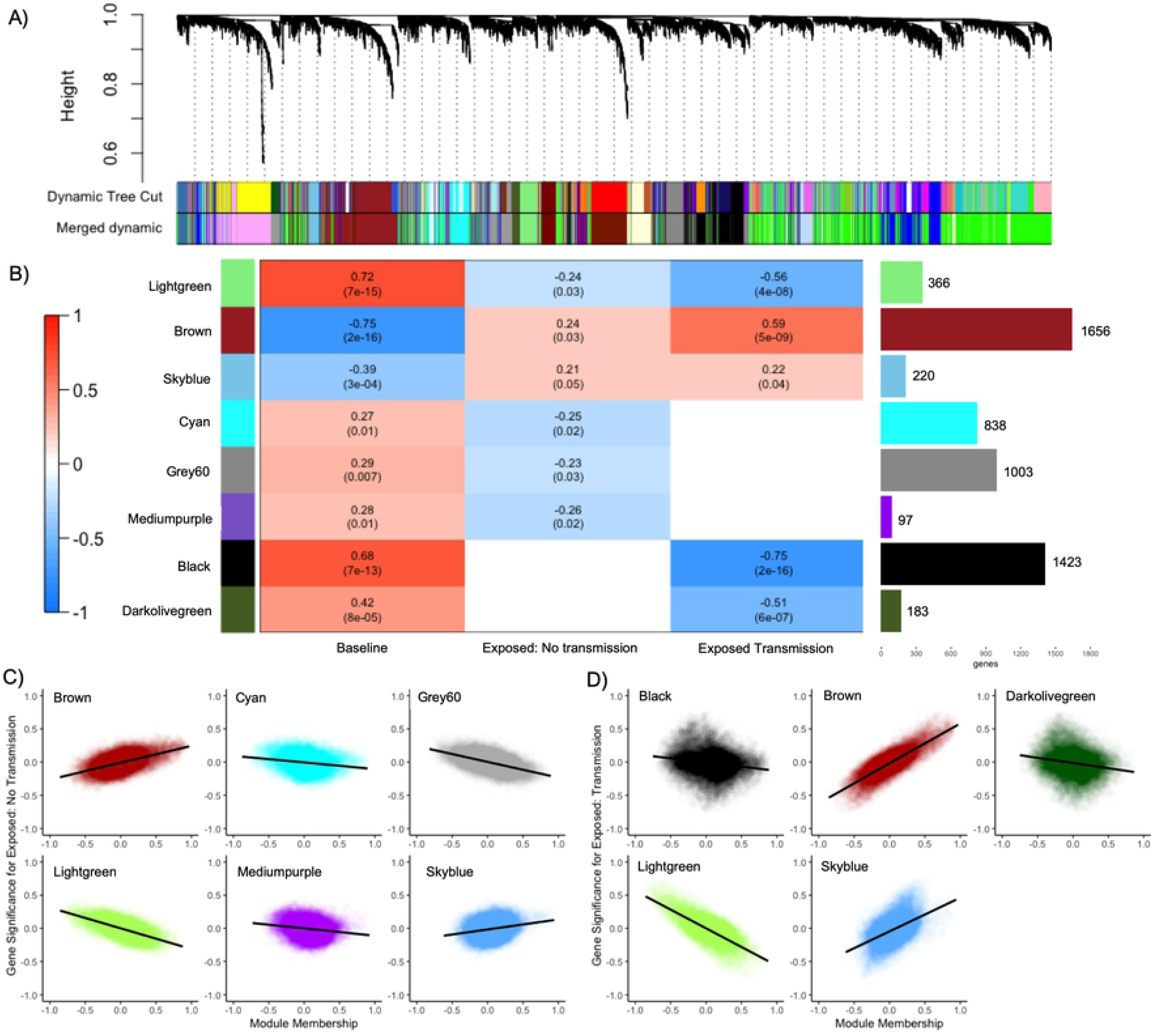
Coexpression analysis identifies 19 gene modules, with eight significantly correlated to Baseline and Exposed corals. A) Dynamic tree height showing merging of modules with similar expression patterns. Merging resulted in the 43 original modules (Dynamic Tree Cut) being merged into 19 modules (Merged dynamic) B) Coexpression heatmap showing the 8 modules that are significantly correlated between Baseline and both disease outcomes (Lightgreen, Brown, Skyblue), Baseline and Exposed: No Transmission (Cyan, Grey60, and Mediumpurple), and Baseline and Exposed: Transmission (Black and Darkolivegreen). Heatmap fill shows positive (red) to negative correlation (blue). The top number in each cell shows the correlation strength, and the bottom number shows module significance to Baseline or disease outcomes. Bar graph to the right shows the number of genes within each module. C) The six modules which are significantly correlated between Baseline and Exposed: No Transmission showing the module membership and gene significance. D) The five modules that are significant between Baseline and Exposed: Transmission showing the module membership and gene significance. For C and D; y-axis shows gene significance which is the absolute value of the correlation between the gene and disease outcome, x-axis shows the module membership which is the correlation of the module eigengene and the gene expression profile.

Of the 19 modules, ‘Lightgreen’ (366 genes, hub gene = Interferon Regulatory Factor 2), ‘Brown’ (1656 genes, hub gene = D-amino-acid oxidase) and ‘Skyblue’ (220 genes, hub gene = PREDICTED: uncharacterized protein LOC107335116) were all significantly correlated (p≤0.05) across Baseline and disease outcomes. These modules were significantly enriched (FDR, p<0.01) for multiple GO Biological Processes, Cellular Components and Molecular Functions with 58 terms for the ‘Brown’ module, 1 for ‘Skyblue’ and 0 for ‘Lightgreen’. The ‘Brown’ module showed a negative correlation for Baseline (R2=-0.75) but was positively correlated for disease outcomes; Exposed: No transmission = 0.24, and Exposed: Transmission = 0.59 (Fig 5, C & D). The ‘Brown’ module was significantly enriched for terms in immune processes such as TLR-6 signaling pathway and MyD88-dependent signaling pathways, positive regulation of cytokine biosynthetic processes, detection and response to diacylated bacterial lipopeptide, podosome, phagocytic and endocytic vesicle membranes, and lipopeptide binding. The ‘Skyblue’ module was significantly enriched for only lipid biosynthetic processes.

Three modules were significantly correlated to Baseline and Exposed: No transmission; ‘Cyan’ (838 genes, hub gene = F-box/LRR-repeat protein 7), ‘Grey60’ (1003 genes, hub gene = Isopentenyl-diphosphate Delta-isomerase 1), and ‘Mediumpurple’ (97 genes, hub gene = pyridoxine-5’-phosphate oxidase) at p≤0.05 (Figure 5, B). The ‘Cyan’ module was significantly enriched (FDR, p<0.01) for 40 GO Biological Processes that included genes involved in cell adhesion, immune responses (complement activation, leukocyte mediated immunity, regulation of coagulation), and metabolic/catabolic processes but showed negative correlations with disease outcomes (Fig 5, C). ‘Mediumpurple’ was enriched for GO terms involved in respiration (electron transport chain, oxidative phosphorylation, ATP synthesis) as well as biosynthetic processes and the positive regulation of necrotic cell death. ‘Grey60’ was enriched for three GO terms: cellular metabolic processes, nitrogen compound metabolic processes and cellular nitrogen compound metabolic processes.

Two modules were significantly correlated to Baseline and Exposed: Transmission; ‘Black’ (1423 genes, hub gene = Ufm1-specific protease 2), and ‘Darkolivegreen’ (183 genes, hub gene = S-adenosylmethionine decarboxylase proenzyme) at p≤0.05 (Figure 5, B). The ‘Black’ module was significantly enriched with one GO term: metabolic processes, while ‘Darkolivegreen’ module was not significantly enriched for any GO terms. All GO terms with associated genes for significant modules are provided in Supp 9.

## Discussion

Our study demonstrates that *A. palmata* mounts a similar immune response to disease as seen in other stony coral species [18–25], with transcriptomic analysis also identifying potential coral and Symbiodiniaceae mechanisms for higher disease virulence in 2017 [30]. We also found that within Symbiodiniaceae, genes linked to ion transport had higher normalized expression in 2017 compared to 2016. This indicates the higher disease prevalence observed by [30] was due to potential dysbiosis with the Symbiodiniaceae.

### Gene expression signatures between 2016 and 2017 show a complementary interaction between Symbiodiniaceae and *A. palmata*

Previously it was found that there were higher rates of disease incidence in the grafting experiments from 2017 compared to 2016 [30]. This increase in disease incidence was hypothesized to be attributed to heightened disease virulence and/or more susceptible genotypes used in 2017 compared to 2016 [30]. In the current study, we included three genotypes that were used in both 2016 and 2017 challenge experiments; HS1, ML6 and CN3 (Table 1). Samples from these genotypes clustered with the year they were exposed to disease, indicating that genotype susceptibility was probably not the cause for the differences in disease prevalence (Fig 2, A). Our results indicate that other factors such as disease type, disease virulence, and the base health of the coral could be potential factors that led to the higher incidence of observed diseases in 2017 [10,26,47]. Additionally, we identified that the *A. palmata* genes driving the difference between 2016 and 2017 were putative coral ribosomal proteins (Fig 3A, Supp 1). These genes had lower overall normalized counts in 2017 compared to 2016 and were involved in translation, rRNA processing, and ribosome biogenesis (Fig 3A). The production of ribosomal proteins are key for the translation of mRNA into proteins, and thus gene expression. The potential reduction in the transcription of these genes in 2017 indicates that the protein production machinery may have been compromised, leading to less physiological and immunological homeostasis and thus higher amounts of disease prevalence [30,48,49].

Conversely, in Symbiodiniaceae, the genes driving the separation of the samples from 2016 and 2017 had higher levels of expression in 2017, with the majority of these genes being involved in ion transmembrane transporter activity (Fig 3B). We hypothesize that this inverse pattern of expression to the *A. palmata* expression could indicate that in 2016 the symbiotic relationship was in equilibrium between the host and Symbiodiniaceae but in 2017, a dysbiosis between host and Symbiodiniaceae was present. This may be due to two potential mechanisms. In 2017, Symbiodiniaceae health may have been limited by certain ions due to a more virulent disease, or an external abiotic factor causing negative impacts. As such, increased expression of genes regulating ion exchange to different molecular compartments was increased to maintain ion balance needed for cellular functions. In Symbiodiniaceae we identified genes encoding for plasma membrane iron permease, voltage-gated sodium channels for sodium transport, ammonium, and nitrate transporters ions. All of these transporters facilitate the movement of iron, ammonium, and nitrate, which are all important for photosynthesis [50–53]. Being limited by these ions causes a lower rate in photosynthetic efficiency and therefore decreases the energy supply provided to the coral host [54, 55]. This may indicate that corals in 2017 had less energy being supplied to the coral host by the Symbiodiniaceae. Alternatively, it has also been shown that in high nutrient environments Symbiodiniaceae within the coral host can become parasitic and can then decrease the translocation of energy to the coral [56]. The Florida Keys have seen increased nutrient loading from anthropogenic sources [57, 58], which has been linked to increases in bleaching and disease susceptibility [59]. This should be tested in the future to fully understand the impacts of higher nutrients on disease susceptibility in *A. palmata* and how it impacts the relationship with the Symbiodiniaceae.

### Enrichment of cell adhesion genes was found in corals that did not show signs of disease

Corals that were exposed to disease, but did not show signs of transmission, had significant differential expression of genes that were enriched for the GO terms: “Cell Adhesion”, and “Cell surface receptor linked signaling pathways” (Supp 4). Cell adhesion is important to maintaining the integrity of the tissue layer and within corals, evidence has been presented that factors, including heat stress and disease, can cause upregulation of genes involved in cell adhesion pathways [20,60–63]. Interestingly, in previous coral disease studies, cell adhesion enrichment was present in corals that were showing signs of disease pathology and hypothesized to be due to the importance of apoptotic processes and phagocytosis of melanized particles and pathogens [20, 21]. Our findings show that cell adhesion is also important in corals not exhibiting visual signs of disease. These processes should be explored further to elucidate differences between corals showing various disease pathologies, and whether these processes are important in genotypic differences in patterns of disease resistance.

### Visual signs of disease are characterized by an innate immune response in A. palmata

PC2 showed a correlation to disease outcome in *A. palmata* (Fig 2B). A common disease response was observed regardless of the year, and this included a number of innate immune processes important in recognition, signaling and effector responses. Most notably genes linked to TLR signaling, the complement cascade, and antimicrobial peptides were present in *A. palmata* exhibiting signs of disease (Fig 4B). These genes may be part of a primary disease response of *A. palmata* and warrant further investigation into their functional significance in overall disease response as well as disease resistance.

*A. palmata* fragments which showed signs of disease had enrichment for GO terms involved in innate immune response including “Defense Response”, “Cytokine Activity”, and “Bioluminescence” (Fig 4B, Supp 5). Our results are similar to previous transcriptomic studies, where innate immunity genes were upregulated in response to disease transmission [18–25]. We identified significantly upregulated TLR 2 and TLR 6 genes which are important innate immune pattern recognition receptors (PRR) that identify gram-negative bacteria and fungi respectively [64, 65]. These receptors are important in initiating the Nuclear Factor Kappa Beta (NF-kB) transcription factor [66–68] that causes production of cytokines and AMPS [69–71]. While other components of the NF-kB pathway were not significantly differentially expressed in this study, they are present in the *A. palmata* genome [34] and have been functionally characterized in the coral *Orbicella faveolata* [68].

Our differential expression results also identified transcripts annotating to AMPs; a bactericidal permeability-increasing protein which has gram-negative bacteria killing properties by targeting the lipopolysaccharide outer layer [72–76], and Achacin, an AMP present in African Giant Slug mucus that has potent gram-positive and gram-negative bacteria killing properties [77–79]. To our knowledge, these AMPs have not been characterized in any other coral disease studies. This is especially notable for *A. cervicornis*, where neither of these have been reported [22, 23]. With the short evolutionary split between *A. palmata* and *A. cervicornis* [80], it would be expected that these AMPs would be present in both species. Re-annotation of past *A. cervicornis* studies with the new *A. cervicornis* genome may identify these AMPs which were previously uncharacterized in the disease response or identify them as unique to *A. palmata*.

Significant differential expression was also identified for 5 lectins including C-type lectin domain family 4 member E and M, Ficolin-1, and Tachylectin-2, and Macrophage mannose receptor 1. These lectins are involved in identifying pathogens and initiating the complement pathway shown to be important in coral symbioses with Symbiodiniaceae [81–83], as well as in response to pathogens and disease [21,24,81,84]. Our findings support previous studies that lectins play a complex role in both symbiosis and pathogen recognition in corals, however, the specific mechanisms and pathways these lectins initiate are still not well understood. A number of genes were also identified to have roles in potential macrophage immune roles. Cationic amino acid transporter has been identified to have a role in macrophage immunity [85], while tyrosine-protein kinase Src42a has been shown to promote macrophages to sites of wounding [86]. Other studies have shown that a sponge has potential macrophage expressed protein activity [87, 88] and that the identified gene is extremely similar to humans identifying a conserved immune process through evolution. While invertebrates do not have adaptive immunity, this may indicate an innate immune phagocytic pathway for managing pathogen infection in *A. palmata*.

### Lipid biosynthesis may play a key role in the activation and maintenance of an immune response in *A. palmata*

The ‘Skyblue’ coexpression module showed a positive correlation with disease outcome (Fig 5B), with significant enrichment of the GO term ‘lipid biosynthetic processes’. This, coupled with the differential gene expression between Baseline and Exposed: Transmission, indicates that *A. palmata* was mounting an energetically expensive immune response to the disease challenge. Furthermore, the enrichment of lipid biosynthetic processes in the ‘skyblue’ module may indicate that the corals sampled in this study have stored energy, in the form of lipids, which can be metabolized and assist in promoting a stronger inflammatory response and fighting off pathogens [89]. This idea has been proposed in other transcriptomic studies on coral disease [25] indicating that this could be integral to multiple coral species disease responses. In the future, linking *A. palmata* lipid production and storage with disease susceptibility may be an important metric for understanding their capacity of resistance and recovery in relation to disease. Genotypes with a higher capacity of lipid production and storage may also be able to initiate a stronger immune response. This has been shown to be true in coral bleaching, namely that individuals with higher lipid stores were able to survive without the Symbiodiniaceae for longer periods of time. This is due to lipids being burnt by the coral resulting in energy allowing continued key life dependent functions [90–92]. This may also be true for disease, with individuals and genotypes with higher lipid stores able to mount a stronger and/or longer immune response.

### ‘Brown’ module is rich in innate immune genes and the hub gene, D-amino acid oxidase, is a critical immune factor involved in *A. palmata* disease response

The “Brown” coexpression module showed increasing positive correlations with disease outcomes (Fig 5, C & D), and included significant enrichment of innate immunity genes and GO terms (Supp 9). Within the “brown” module, D-amino acid oxidase (DAO) was identified as the hub gene. DAO is a peroxisomal enzyme that has been identified to be important in mucosal microbiome homeostasis and leukocyte phagocytic activity in mammalian models [93–96]. In corals this enzyme, to our knowledge, has not been documented in response to disease, and its presence as a hub gene in disease response could indicate that it is a critical immune factor that has previously been overlooked. In mammalian models free-floating D-amino acids (DAA), which are actively released by bacteria, are catalyzed by DAO. Phagocytic cells have been shown to be chemo attracted to free-floating DAA by recognition through G-coupled protein receptors [93]. There are a number of G-coupled protein receptors present in the “brown” module (Supp 8), indicating these may be involved in the recognition of DAA during phagocytosis in *A. palmata*. During bacterial phagocytosis, DAO is released into the phagosome, catalyzing the deanimination of DAA which releases hydrogen peroxide and kills the bacteria. This enzyme may also have greater implications in coral host-microbiome interactions. Beneficial holobiont bacteria have been shown to have resistance to host DAO while also being able to manage levels through the TLR-to-NF-kappa-B pathway [95]. We therefore hypothesize that DAO could have a dual role in *A. palmata* as it is important in the immune response, as well as maintaining symbiosis with coral microbial partners as in other organisms [95] with future research needed to characterize its role.

## Conclusions and future directions

Within this study, we present evidence that *A*. *palmata* initiated an immune response to a disease challenge assay. We identified genes linked to corals that showed no signs of disease (Exposed: No transmission) which is a factor to consider when looking at disease resistance. Furthermore, we identified sets of genes that show high similarity to other coral disease transcriptomic studies [18–25] when disease signs are visually present (Exposed: Transmission). Since the response to disease exposure in *A. palmata* is similar to other coral species, we can now start to explore putative genetic mechanisms which confer genotypic disease resistance, and mechanisms that could cause increases in disease susceptibility. This work has important implications for restoration practitioners, as it may help increase outplant survival efforts through development of novel diagnostic markers and identification of genotypic resistance, while also expanding the current knowledge on the evolutionary history of innate immunity in corals and invertebrates.

## Acknowledgements

We would like to acknowledge the team of field workers who contributed to the field challenge disease assays which included: Allan Bright, Rachel Pausch, Annie Peterson, Emma

Pontes, and Phil Colburn. We also like to thank the coral nurseries [Coral Restoration Foundation, Florida Fish and Wildlife Commission, and Dr. Lirman (University of Miami)] for providing the coral fragments and CRF for permitting work to be conducted in their nursery.

## Author contributions

X.M.S., M.W.M., and D.W. designed and performed the field challenge disease assays. B.D.Y, X.M.S AND N.T.K identified samples to sequence for transcriptomic analysis. B.D.Y and X.M.S performed laboratory work. B.D.Y performed all bioinformatic analysis, figure and table preparation, B.D.Y, S.R and N.T.K performed manuscript writing. All authors reviewed drafts of the paper before submission.

## Supporting Information

**Supp 1**: *A. palmata* Principal Component 1 gene loadings and GO list with associated genes.

**Supp 2: Heatmap of *Symbiodiniaceae* genes associated with photosynthetic GO terms.**

**Supp 3**: ***Symbiodiniaceae* Principal Component 1 gene loadings and GO list with associated genes.** For left heatmap grey = 2016 samples, black = 2017 samples. Fill shows higher (red) to low (blue) gene counts using a variance stabilizing transformation. Column dendrogram shows hierarchical clustering of samples. Rows (genes) also arranged using hierarchical clustering with dendrogram omitted. Right heatmap is presence (black) and absence (white) of genes to GO terms.

**Supp 4: Baseline VS Exposed: No Transmission DeSeq2 results and significant GO terms with associated genes.**

**Supp 5: Baseline VS Exposed: Transmission DeSeq2 results and significant GO terms with associated genes.**

**Supp 6: Shared genes between DeSeq2 contrasts.**

**Supp 7**: **Coexpression heatmap for the 19 modules identified from WGCNA analysis.** Heatmap fill shows positive (red) to negative correlation (blue). The top number in each cell shows the correlation strength, and the bottom number shows module significance to Baseline or disease outcomes.

**Supp 8: Gene lists for significant modules from WGCNA analysis.**

**Supp 9: GO terms and associated genes for significant WGCNA modules.**

